# Heimdallarchaea encodes profilin with eukaryotic-like actin regulation and polyproline binding

**DOI:** 10.1101/2020.04.23.051748

**Authors:** Sabeen Survery, Fredrik Hurtig, Syed Razaul Haq, Ann-Christin Lindås, Celestine N. Chi

## Abstract

The evolutionary events which led to the first eukaryotic cell are still controversial^1-4^. The Asgard genome encodes a variety of eukaryotic signature proteins previously unseen in prokaryotes. Functional and structural characterization of these proteins is beginning to shed light on the complexity and pedigree of the ancestral eukaryotic cell^5,6^. In eukaryotes, the key cytoskeletal protein actin is important for diverse cellular processes such as membrane remodeling and cell motility^7^. Dynamic polymerization of actin provides both structure and generates the force which drives motility and membrane remodeling. These processes demand rapid filament assembly and disassembly on microsecond timescales. In eukaryotes, a variety of highly adapted proteins including gelsolin, profilin, VASP, ARP2/3 and signaling molecules (Phosphatidylinositol-4,5-bisphosphate (PIP_2_)) are crucial for organizing cellular cytoskeleton dynamics. Amongst others, the Asgard genomes encode predicted putative profilin homologues that regulate eukaryotic actin polymerization *in vitro*^5,8^. Interestingly, Asgard profilins appear to be regulated by PIP_2_, but not by polyproline rich motifs which are important for recruitment of actin:profilin complexes in eukaryotes^5,9^. These findings indicate that the Asgard archaea may have possessed analogous membrane organization to present-day eukaryotes, but that polyproline-mediated profilin regulation may have emerge later in the eukaryotic lineage^5^. Here, we show that Heimdallarchaeota, a candidate phylum within the Asgard superphylum, encodes a putative profilin (heimProfilin) that interacts with PIP_2_ and is regulated by polyproline motifs, implicative of an origin predating the rise of the eukaryotes. Additionally, we provide evidence for a novel regulatory mechanism whereby an extended N-terminal loop abolishes PIP_2_ and polyproline interactions. Lastly, we provide the first evidence for actin polymerization of an Asgard actin homologue. In context, these findings provide further evidence for the existence of a complex cytoskeleton already in Last eukaryotic common ancestor (LECA).

## Results and Discussion

The recent discovery of the Asgard superphylum represents a major breakthrough in the study of eukaryogenesis^1,8^. While the Asgard phyla are predicted to encode a large number of eukaryotic signature proteins (ESPs), only a limited knowledge is available on the actual eukaryotic-like function of these Asgards genes^5,10^. To verify that Heimdallarchaeota encode a *bona fide* profilin, we determined the 3D protein structure using nuclear magnetic resonance (NMR). Several profilin structures from the Asgard superphylum including Loki profilin-1, Loki profilin-2 and Odin profilin have been determined previously by X-Ray crystallography both individually and bound to rabbit actin^5^. However, there are considerable phylogenetic differences separating the known Asgard phyla, and Heimdallarchaeota is currently thought to be more closely related to eukaryotes than any other Asgard phyla^8^. Nevertheless, sequence conservation amongst the Asgard profilin homologues is relatively low and identity is mostly established through structural homology. At a first glance, our NMR structure depicts a typical profilin fold, with seven strands interlinked by loops connecting four helices (Fig. 1). However, the orientation, positions and length of the helices and loops differ dramatically compared with Loki profilin-1 and canonical eukaryotic profilins. Detailed structural comparison reveals that Heimdallarchaeota profilin (heimProfilin) is divergent from the Loki profilin-1 (root mean squared deviation (RMSD) > 3.75 Å (Supplementary Fig. 1). Notably, differences include the formation of an additional helix between residues H123-S129, the re-orientation of the N-terminal helix (residues S27-Q35) to an open conformation, a shorter Loki-loop, the absence of a helix between residues G72-P75 and the presence of a long disordered N-terminal loop (residues 1-20) (Fig. 1c). These structural differences indicate that despite the overall profilin fold, the heimProfilin differs to the eukaryotic and the recently determined Loki profilins.

**Figure 1.**
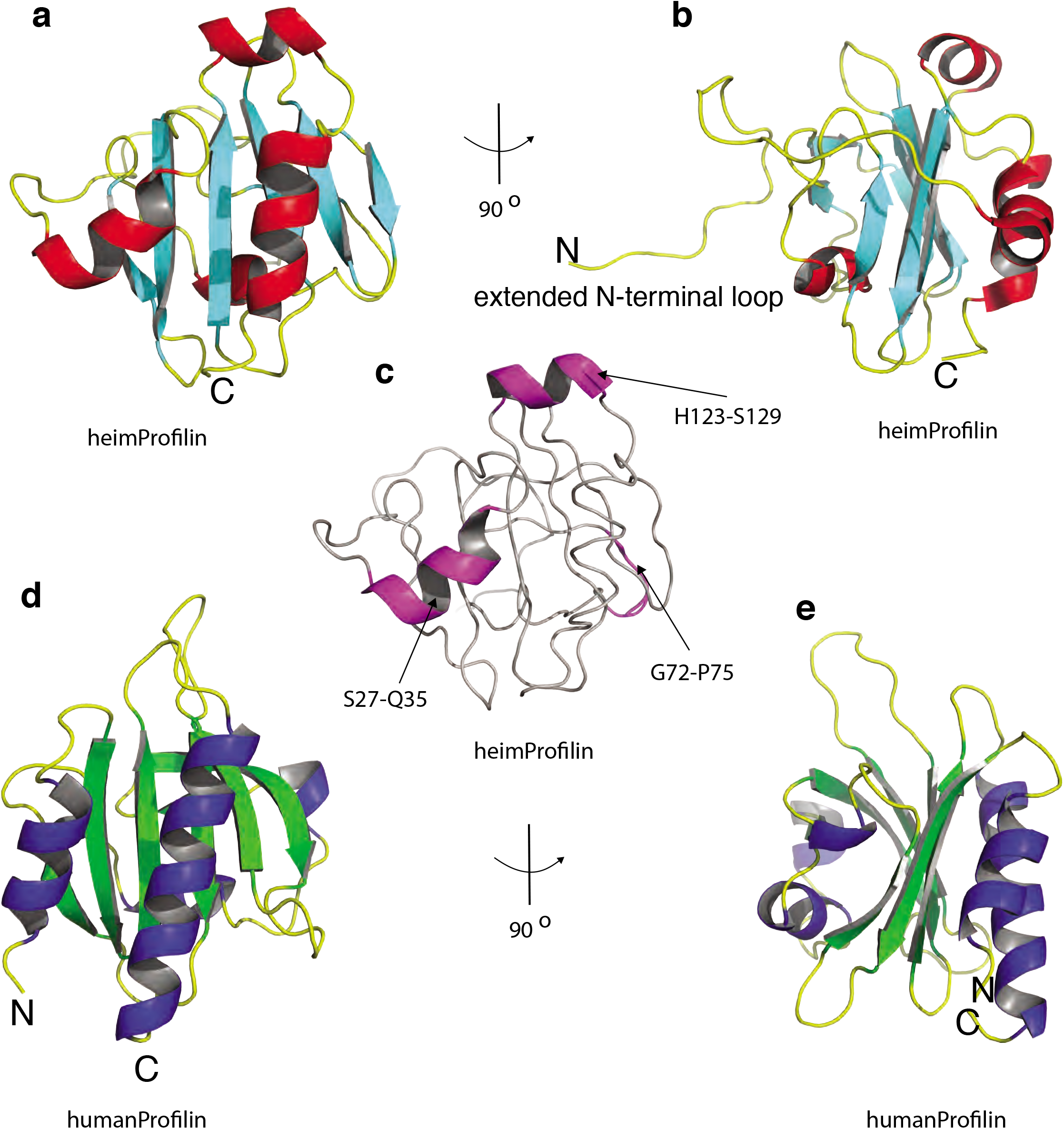
Heimdallarchaeota encodes profilin with extended structures. **a**, Schematic of the structure of heimProfilin. The orientations of the N- and C-terminal helices are displayed. In addition, the helix between residues H123-S129 is also shown. **b**, Reorientation of the structure in a by 90° to show the extended N-terminal loop between residues 1-124. **c**, Schematic of heimProfilin showing notable differences in structural elements to that of human profilin **d** and **e**, Schematics of human profilin-1 are also displayed for comparison. Note that the N-terminal helix in heimProfilin is reoriented by almost 60° as compared to the human profilin-1. The human profilin does not harbor the N-terminal extension and the N-terminal helix is slightly longer.

The extended N-terminal loop in heimProfilin is completely absent in Loki profilin and eukaryotic profilin. To investigate the loop’s function, we cloned and expressed a truncated form of heimProfilin which we called ΔN-heimProfilin which lacked the extended N-terminal loop (residues 1-23). The overall fold of the ΔN-heimProfilin was similar to that of the heimProfilin as judged from NMR backbone ^15^N-^1^H and ^13^Cα chemical shifts (Supplementary Fig. 2). To further investigate the functions of both profilins, we allowed rabbit actin to polymerize in the presence of heimProfilin or ΔN-heimProfilin and observed the resulting filament network with Airyscan super-resolution microscopy. In these experiments heimProfilin was able to inhibit filament network formation in a concentration dependent manner (Fig. 2a). In contrast, ΔN-heimProfilin did not alter the filament network (Fig. 2a). To verify these results, we followed the polymerization dynamics of pyrene labeled rabbit muscle actin in the presence of heimProfilin or ΔN-heimProfilin. In line with the microscopy data, we found that heimProfilin was able to modulate rabbit actin polymerization in a concentration dependent manner (Fig. 2b). Conversely, ΔN-heimProfilin did not alter rabbit actin polymerization (Fig. 2c). However, sedimentation assay data showed that both heimProfilin and ΔN-heimProfilin were able to interact with rabbit actin (Supplementary Fig. 3d, e). These results indicate that the N-terminal loop is essential for regulation of actin polymerization dynamics, but not for actin binding and is indicative of the functional role for the extended N-terminal loop previously unseen in eukaryotic or Asgard profilin homologues. To further verify these results, we first cloned, expressed and purified an actin homologue from Heimdallarchaeota. We used two variants of Heimdall actin; a full length (heimActin) and a truncation mutant form (ΔC-lieimΛctin) where the last 35 C-terminal amino acids, crucial for polymerization, had been deleted^7,11^. Electron microscopy showed that purified heimActin could form thin filaments while the truncation mutant could not (Fig. 2d) and that the heimActin was more active in ATPase assay than the ΔC-heimActin (Fig. 2e). We then compared the sedimentation profiles of the heimActin and ΔC-heimActin in presence of heimProfilin and ΔN-heimProfilin. While both heimProfilin and ΔN-heimProfilin were seen to interact with heimActin and ΔC-heimActin respectively, we observed that only heimProfilin was able to modulate the polymerization of heimActin (Fig. 2f-h) corroborating the above polymerization modulation of rabbit actin by heimProfilin. This finding indicates that the Heimdallarchaeota possess profilins which are able to regulate both heimActin as well as eukaryotic actin polymerization.

**Figure 2.**
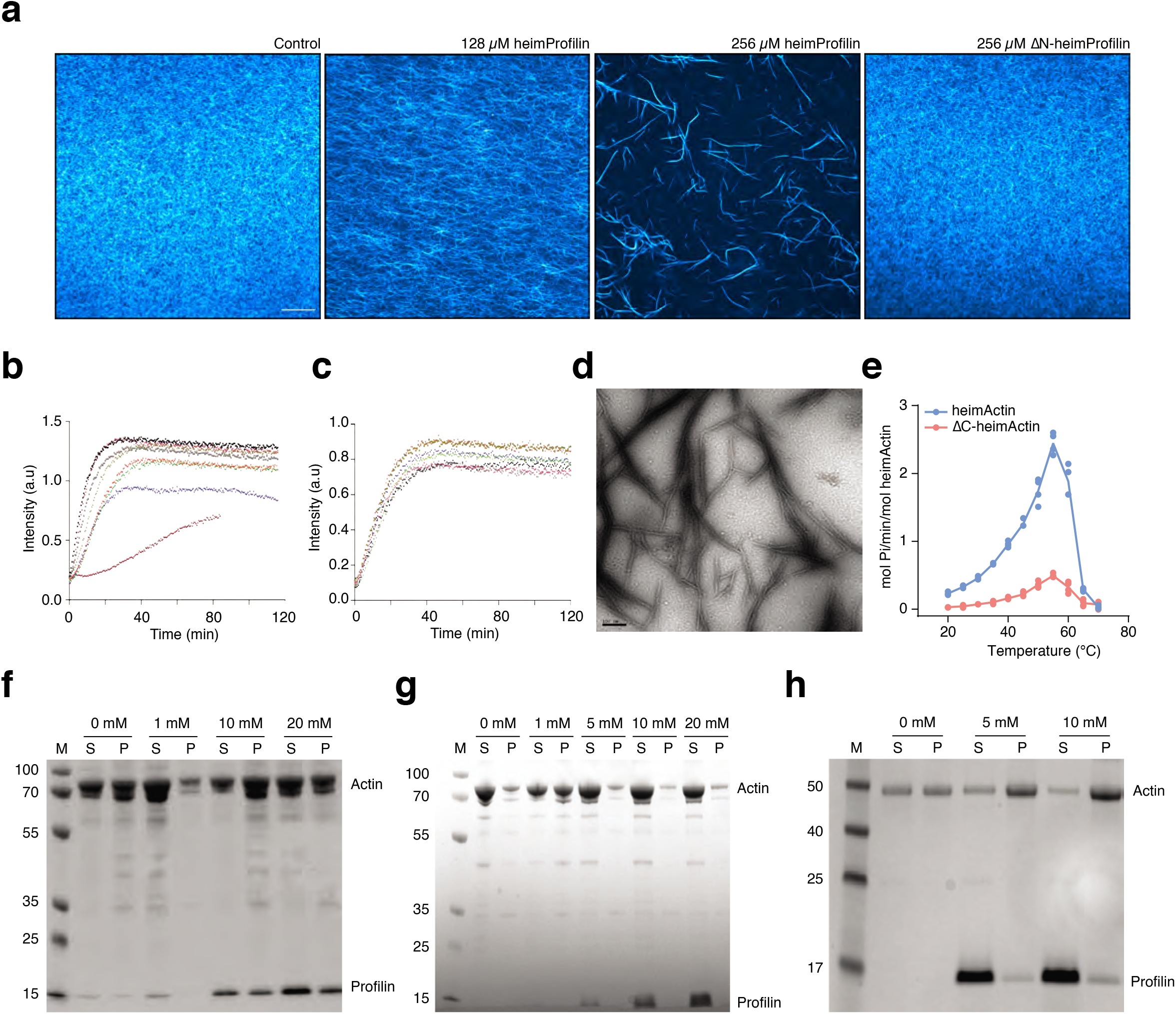
Heimdallarchaeota encodes actin (heimActin) that polymerizes and is modulated by its heimProfilin. **a,** Airyscan super-resolution microscopy of rabbit actin in presence of different concentrations of heimProfilin; 0 μM, 128 μM and 256 μM, or 256 μM of NΔ-heimProfilin**. b,** Pyrene-polymerization profiles of 2 μM rabbit actin (10% pyrene-labeled) alone (pink) or with different concentrations of heimProfilin; 19 μM (black), 48 μM (grey), 93 μM (olive), 137 μM (red), 200 μM (green), 250 μM (blue) and 280μM (magenta). **c,** Pyrene-polymerization profiles of 1 μM of rabbit actin (10% pyrene-labeled) alone (pink) or with different concentrations of ΔN-heimProfilin; 20 μM (black), 50 μM (grey), 100 μM (olive), 150 μM (red), 200 μM (green), 250 μM (blue). **d,** Electron microscopy (EM) images of heimActin forming thin, uniform filamentous polymers. **e**, Phosphate released during polymerization of heimActin (blue) or ΔC-heimActin (red) as a function of temperature. **f,** Sedimentation assay for heimActin alone and with increasing concentrations of heimProfilin. **g**, Same as in f) but with heimActin and ΔN-heimProfilin. Here, both ΔN-heimProfilin and heimActin appear in the soluble fraction but not in the pellet, indicating that ΔN-heimProfilin do not increase the polymerization of heimActin. **h**, Presence of heimProfilin increases the polymerization of ΔC-heimActin.

To further investigate the interaction between heimProfilin and heimActin and to see which residues are required for the interaction, we turned to nuclear magnetic resonance (NMR) spectroscopy and performed binding titrations between heimProfilin, heimActin and their respective mutants. We observed chemical shift changes for the interaction between both heimProfilin and ΔN-heimProfilin with heimActin, in line with the pyrene polymerization and sedimentation assays (Fig. 2 and Supplementary Fig. 4). ΔN-heimProfilin exhibited stronger chemical shift changes than heimProfilin in the presence of heimActin when comparing similar amino acid residues at similar concentrations, indicating the N-terminal loop might modulate binding in a manner unseen in the sedimentation assays. We also observed that the interaction between ΔN-heimProfilin and ΔC-heimActin was modest (few residues exhibiting small chemical shift changes) in line with the sedimentation assay (Supplementary Fig. 5). From the NMR titrations, we were able to map the site of interaction (Supplementary Fig. 1 and 4), which corresponded to the following residues in heimProfilin: N31, W59 and W59 side-chain, S64, Q69, W70, F84, G103, G104, I106, N113, T127, E139. For binding experiments with ΔC-heimActin, we observed chemical shift changes for the following residues in ΔN-heimProfilin: Y26, Y41, I51, W59, G63, Q69, M71, G104, N113, G130. For comparison, the residues responsible for actin interaction in eukaryotes are shown on the sequence alignment for both archaeal and eukaryotic profilin (Supplementary Fig. 1). Together these results indicate that the Heimdallarchaeota profilin is functional and is consistent with dynamic barbed end binding of actin.

Polyproline binding from the enabled/vasodilator-stimulated phosphoprotein (Ena/VASP) family of proteins is important for nucleation and elongation of actin filaments^12^. To verify if heimProfilin binds to polyproline we performed binding experiments both by NMR spectroscopy and Isothermal titration calorimetry (ITC), using heimProfilin and ΔN-heimProfilin. We observed a moderate binding of ΔN-heimProfilin to polyproline with an affinity constant of ∼ 200 μM and a very weak binding for the heimProfilin with affinity constant in the higher micro molar range (Fig. 3). Titration by NMR reveals that the residues responsible for polyproline binding were K22, G49, Y52, W53 and W53 side-chain, I106, A111, A145, F147 and Q148 (Supplementary Fig. 1). Revisiting the structure of heimProfilin and comparing it with that of eukaryotic profilin reveals that the N-terminal helix is orientated upwards creating a pocket which allows the polyproline motif to bind in an “L-like” fashion as compared to the human profilin-polyproline binding (Supplementary Fig. 1). These striking observations explains the reason why Loki profilin-1, −2, and Odin profilins could not interact with polyproline motifs, whereas heimProfilin could. Structural data from Loki profilin-1, −2, and Odin profilins indicated that their N-and C-terminal helices parallel and are more close to each other making this type of interaction highly unlikely^5^. These results suggest that, contrarily to what was previously thought, polyproline binding (to profilin) could have emerged before the split between the Asgard and eukaryotic lineages. This pose the question, why do profilin from Loki and Odin do not possess the N-terminal loop extension? One explanation could be that some Asgards lost this loop, or conversely, that Heimdallarchaea acquired the loop independently by convergent evolution, or through horizontal gene transfer. We performed a blast search and alignment analysis of all profilins deposited in NCBI in an attempt to identify any indication of additional structural elements upstream from the known start position of all other profilins to verify that the genes were correctly annotated. Interestingly, we observed that several profilins, mainly from the Thorarchaeota (TF12995.1, TFG09823.1, TFG30347.1, TFF94849.1, RLI55859.1), contained sequences with N-terminal extension ranging between 5 and 22 residues, upstream of their previous designated start position (Supplementary Fig. 6). In attempt to see if these thorProfilin retain the profilin fold and if they also possesses the N-terminal loop, we modeled the 3D structure for one of the thorProfilin, TF12995.1, which contains 22 amino acids upstream the known start position using the online software RaptorX^13^. Indeed, we found that the overall fold of the predicted structure matches our 3D structure determined for heimProfilin with an additional N-terminal extension (Supplementary Fig. 7). This indicates that these N-terminal extensions are present in other archaea and could potentially play similar roles as observed for heimProfilin albeit maybe not to the same extent.

**Figure 3.**
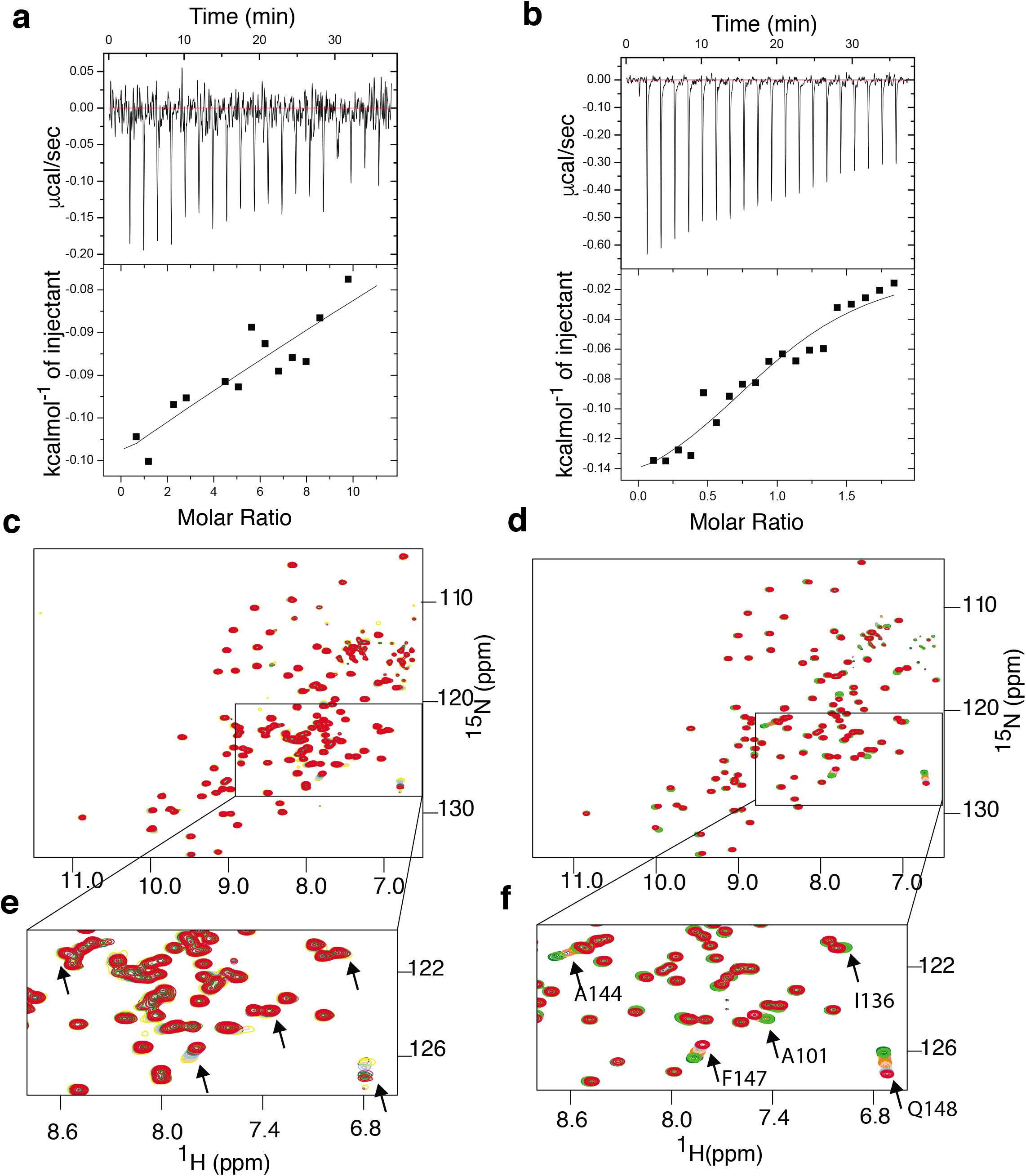
Heimdallarchaeota profilin interacts with polyproline. Isothermal titration calorimetric (ITC) binding measurements between heimProfilin **a)** or ΔN-heimProfilin **b)** with polyproline motifs of VASP (PPPAPPLPAAQ). The heimProfilin showed weaker binding strength compared to the ΔN-heimProfilin which had a *K*_D_ of ∼200μM. **c and d,** Nuclear magnetic resonance (NMR) ^1^H-^15^N chemical shifts for the binding reaction of heimProfilin and ΔN-heimProfilin with polyproline (VASP) respectively. **e** and **f,** Expansions from c) and d) showing the movement of some interacting residues as the concentration of polyproline increases.

In eukaryotes membrane phospholipids, particularly Phosphatidylinositol-4,5-bisphosphate (PIP_2_), regulate the activities of many actin binding proteins including profilin, cofilin, ezrin, Dia2, N-WASP and meosin^14^. It should be noted that Asgard archaea likely do not possess similar eukaryotic membrane architecture. However, they do express membranes with lipids that have similar features. For example, some archaea lipids have similar inositol head groups but varied tails known as archaeols^15^. Therefore, we verified whether heimProfilin is able to interact with PIP_2_ by monitoring changes in NMR chemical shifts upon addition of PIP_2_ into a solution of heimProfilin or ΔN-heimProfilin. We observed that while ΔN-heimProfilin interacted with PIP_2_, no interaction was observed for the heimProfilin (Fig. 4). The residues responsible for phospholipid binding in the Asgard superphyla have not been mapped before and were only speculated from surface charge distribution^5^. These NMR titration experiments gave us a perfect opportunity to mapped this binding site. The following residues were observed to display chemical shift perturbation upon addition of PIP_2_: S27, D28, L30, N31, Q35, S36, V43, G49, N99, K110, A111, F117, L118, S119, E139, I140, M142, M143, K146, F147, Q148. Although, the change in chemical shift of specific residue does not mean a direct interaction of that residue, we observed that all the residues displaying large changes in chemical shift were located on the same surface of the protein (Fig. 4). This strongly indicates the binding interphase for PIP_2_. We also investigated the potential interaction of inositol trisphosphate (IP_3_), a second messenger signaling molecule resulting from the hydrolysis of PIP_2_. However, we did not observe any interaction for IP_3_ with either heimProfilin or the ΔN-heimProfilin (Fig. 4e, f and h). Taken together, these observations indicate that, i) Asgard profilins interact with phospholipids, implicating phospholipids in Asgard archaea actin modulation and ii) profilin from Heimdallarchaeota possess an extended N-terminal loop involved in actin polymerization regulation (Supplementary Fig. 8a). Interestingly, deleting this loop enhanced binding to both PIP_2_ and polyproline motifs; two important partners in modulation of actin cytoskeleton dynamics in eukaryotes. In addition, we observed from sequence analysis that other profilins possess potential N-terminal extensions that could play a role in modulating actin polymerization, indicating that polyproline-mediated regulation could predate the Asgard-Eukarya split. We propose a model where modification of the extended N-terminal loop, or interaction with third-party proteins, causes the heimProfilin protein to behave similarly to ΔN-heimProfilin. This inhibits modulation of actin polymerization dynamics but allows for PIP_2_ and polyproline interactions. Concurring or subsequent demodification would flip heimProfilin to an actin modulating state, allowing for actin polymerization regulation. Modification or interaction with third-party proteins would then be able to reset profilin to the first step of the cycle (Supplementary Fig. 8b). In conclusion, this study suggests that Asgard archaea encode a complex cytoskeleton functionally analogous to major eukaryotic cytoskeletal characteristics. Moreover, Heimdallarchaeota expresses profilins that are potentially regulated by phospholipid binding and polyproline interaction, something which was long thought to be eukaryotic-specific, and previously not observed in other Asgard archaea.

**Figure 4.**
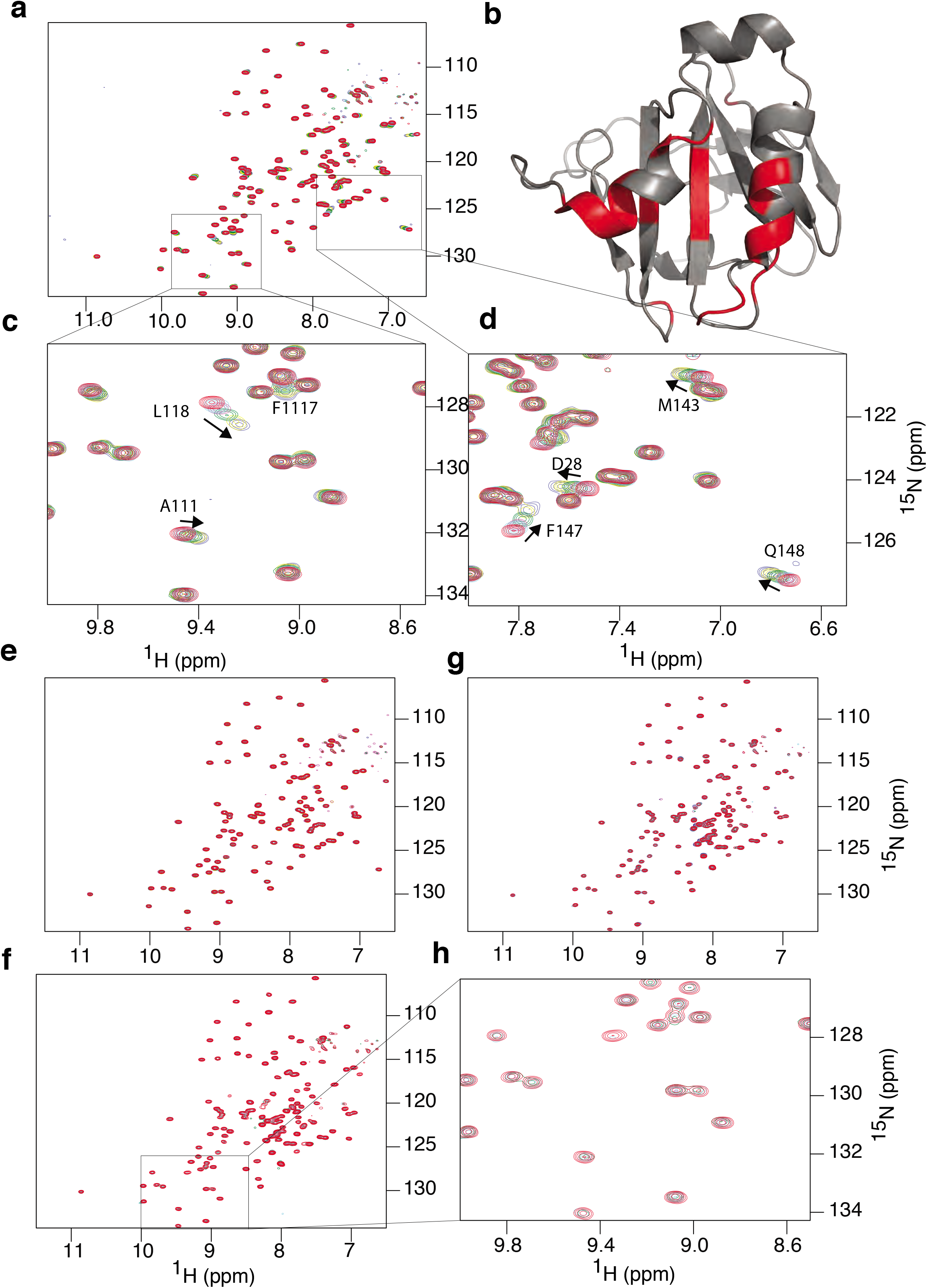
HeimProfilin N-terminal extension is important for interaction with phospholipids. **a,** Overlay ^1^H-^15^N TROSY-HSQCs of ΔN-heimProfilin (400 μM) with increasing concentrations of Phosphatidylinositol-4,5-biphosphate (PIP_2_); 0 μM (red) 150 μM (cyan), 300 μM (magenta), 600 μM (green), 1200 μM (yellow), 2100 μM (blue). **b,** Schematic of heimProfilin with the interacting residues color coded. The interaction site appears to be located between the N- and C-terminal helix. **c-d**, Expansion of a) showing a few residues belonging to the b-strand residues F117-W120 and residues Q110-A111 are also strongly involved in the interaction. **e**, Overlay ^1^H-^15^N TROSY-HSQCs of ΔN-heimProfilin (300 μM) with increasing concentrations of D-myo-inositol-1,4,5-triphosphate (IP_3_); 0 μM (red) 150 μM (cyan), 300 μM (magenta), 600 μM (green), 1200 μM (yellow), 2100 μM (blue). **f**, Similar to a) but with heimProfilin and PIP_2_. **g**, Similar to e) but with heimProfilin interacting with IP_3_. No chemical shift changes were observed for heimProfilin-IP3 interaction. **h**, Expansion of a region in f).

## Author’s contribution

S. S, F. H, S. R. H, A-C. L and C. C conceived the project. S. S, S. R. H and FH cloned, expressed and purified all proteins. SS and SRH performed the pyrene and sedimentation assay. FH performed the ATP assay and confocal microscopy. S. S performed the electron microscopy. C. C performed all NMR and ITC experiments. C. C wrote the paper with contributions from all other authors.

## Ethics approval

not applicable.

## Consent to participate

not applicable.

## Consent for publication

all authors read and approved the manuscript.

## Availability of data and material

all data and material are available and can be obtain from the authors

