## supplemental info for "Heimdallarchaea encodes profilin with eukaryotic-like actin regulation and polyproline binding"

### Materials and Methods

#### Protein expression and purification

The profilin (sequence ID OLS22855.1) and actin (sequence ID OLS30618.1 and TET76256.1, respectively) used in this study were from Heimdallarchaeota. Sequence OLS30618.1 has a 35 amino acids C-terminal deletion compared with TET76256.1. Thus the OLS30618.1 variant is from now on named ΔC-heimActin and the TET76256.1 heimActin. Both genes were sub cloned into the pSUMO-YHRC vector (kindly provided by Claes Andréasson (Addgene Plasmid #54336; RRID: Addgene_54336)) with an N-terminal 6xHistidine-tag and a SUMO-tag (cleavable with Ulp1 protease). Recombinant proteins were overexpressed in *E. coli* Rosetta (DE3) strain. Initially, the cells were grown at 37 °C in 2 x TY broth. Protein expression was induced with 0.5 mM IPTG when the optical density at $A_{600}$ was 0.6 - 0.8. After induction, the cells were grown overnight at 30 °C. For Actin expression, the cells were grown for 4 h at 20 °C post induction. The cells were harvested by centrifugation and the cell pellet was dissolved in binding buffer; 50 mM Tris-HCl pH 8.0 or pH7.5 (for heimProfilin), 0.3 M NaCl/KCl, 1 mM TCEP, 10 mM imidazole, 10% glycerol. For actin, the buffer was supplemented with 2 mM $MgCl_2$.

#### Protein Expression for NMR

The pSUMO-YHRC vector with the heimProfilin (sequence ID OLS22855.1) and ΔN-heimProfilin (with the first 1-23 amino acids in the N-terminal loop deleted from the sequence OLS22855.1) was transformed and expressed in *E. coli* Rosetta DE 3 cells. The cells were grown in 2 x TY media at 37 °C until $A_{600}$ = 0.8. The cells were harvested by centrifugation at 4,000 x g for 15 min and washed twice with M9 medium. The cells were thereafter grown overnight at 30 °C in M9 medium, supplemented with 1g/L $^{15}$N-ammonium chloride and 1g/L $^{13}$C- glucose. Protein expression was induced with 0.5 mM IPTG. For the labeling of protein with Deuterium ($^{2}$H), the M9 medium was made in $D_2O$. The cells were harvested by centrifugation and the cell pellet was dissolved in the binding buffer; 50 mM Tris-HCl pH 7.5, 0.3 M NaCl, 1 mM TCEP, 10 mM imidazole, 10% glycerol. Cells were lysed by sonication, followed by centrifugation at 25,000 x g for 45 min. The supernatant was loaded onto His-GraviTrap column (1ml, GE healthcare) pre-equilibrated with binding buffer. The 6xhistidine tagged proteins bound to the column were eluted with binding buffer containing 250 mM Imidazole. The eluted proteins were incubated with ULP1 protease overnight at 4 °C for tag cleavage, followed by buffer exchange on PD-10 (GE

Healthcare). The tag was removed by reloading the protein solution onto the His-GraviTrap column. The proteins were concentrated using a 10,000 NMWL cutoff centrifugal filter (Merck-Millipore). The concentrated proteins were subjected to size-exclusion chromatography on Superdex-200 or Superdex-75, 10/300 GL column (GE healthcare) as the final purification step. For size-exclusion chromatography, the column was pre-equilibrated with 25 mM Tris-HCl pH 8.0 (or 7.5 for profilin), 50 mM NaCl, 1 mM TCEP, 1 mM $MgCl_2$ (for actin only) and 10% glycerol. Fractions containing the purified proteins were pooled, concentrated, and stored at -80 °C for further use.

**Nuclear magnetic resonance experiments**

All NMR experiments were done on Bruker spectrometers equipped with tripled resonance cryogenic probes operating at proton larmor frequencies of 600, 700 and 800 MHz. Experiments for assignment were as previously reported[1]. All NMR binding titrations were done on the Bruker 600 MHz at 298K unless otherwise stated. All protein samples were either single labeled $^{15}N$, or double labeled $^{15}N$, $^{13}C$, at concentrations between 5-10 mg/ml in 25 mM Tris-HCl pH 7.5, 50 mM NaCl, 5% glycerol and supplemented with 3% $D_2O$ and 0.03% sodium azide. 2D $^{1}H$-$^{15}N$ TROSY-HSQC experiments were recorded at a fixed amount of profilin or NΔ-heimProfilin (200-400 μM) with increasing amounts of heimActin, ΔC-heimActin, polyproline (PPPAPPLPAAQ), or phospholipids (Phosphatidylinositol-4,5-bisphosphate (PtdIns (4,5)$P_2$ ($PIP_2$) and of D-myo-inositol-1,4,5-triphosphate ($IP_3$)). Experiments involving heimActin or ΔC-heimActin were done in the presence of 0.9 mM latrunculin to inhibit polymerization. All experiments were processed with Bruker TopSpin software and analyzed with the CcpNmr analysis program[2].

**Structure determination**

The assignments of Heimdallarchaeota profilin was as described[1]. In addition, we measured 3D $^{1}H$-$^{1}H$ NOESY resolved in $^{13}C$-$^{1}H$ and $^{15}N$-$^{1}H$ TROSY experiments with the following specifications: 80 ms mixing time and 128 ($^{15}N$ or $^{13}C$) × 256 ($^{1}H$) × 2048 ($^{1}H$, direct) and used for distant restraint determination. We also used the $^{3}J_{HNH}\alpha$ couplings and together with the NOE derived distances were deposited in BMRB with ID: 50190. Structure calculations were done using the CYANA 3.98.13[3] package in two steps. First, the NOESY cross peaks were converted into upper distance restraints in an automated process in CYANA. The distance restraints in addition to the $\phi/\psi$ dihedral angles determined from Carbon α-chemical shifts and $^{3}J_{HNH}\alpha$ were used as input for the initial structure calculations. The structures were calculated with 200,000 torsion angle dynamics steps for 100 conformers starting from random torsion angles by simulated annealing. For representation and analysis, the 20 conformers with the lowest target function values were selected. The structural statistics together with all input data for the structure calculations are presented in supplementary Table 1. The structural coordinates have been deposited in the protein data bank with PDB code: 6YRR

**Electron Microscopy**

For EM observation, heimActin (5 μM) was polymerized for 2 h and ultracentrifuge for 1 h at 150,000 x g at 4 °C. The pellet was dissolved in F-actin buffer (20 mM Tris-HCl pH 8.0, 200 mM KCl, 1 mM ATP, 4 mM $MgCl_2$) and applied to carbon-coated grids for 60 s and negatively stained with 1% (w/v) uranyl acetate. A TECNAI G2 spirit Bio-TWIN electron microscope (FEI company) was used at an accelerating voltage of 80 kV with a 70-μm objective aperture and a 100-μm condenser aperture at a nominal magnification of 1.8-3.0 X $10^4$.

**Confocal Laser Scanning Microscopy:** Coverslips were thoroughly cleaned with ethanol before covering with 2% poly-L-lysine (ThermoFisher) for 1 h at 37 °C. Coated coverslips were washed once with distilled water before air-drying. Rabbit actin (5uM) in G-buffer (5 mM Tris-HCl pH 7.5, 0.5 mM DTT, 0.1 mM $CaCl_2$, 0.5mM ATP). Actin was polymerized by adding 100 mM KCl/1 mM $MgCl_2$ supplemented with heimProfilin or ΔN-heimProfilin at room temperature for 40 min. Polymerized actin was then gently diluted to 1 μM with actin polymerization buffer (10 mM Tris-HCl pH 7.5, 50 mM KCl, 2 mM $MgCl_2$) and supplemented with 1 μM Phalloidin-FITC (Sigma) for visualization. Of this solution, 15 μl was gently aspirated onto a cleaned microscopy slide before the addition of a coated coverslip. Imaging was executed using a Zeiss LSM 800 Airyscan using a Plan-Apo 63x/1.40 Oil DIC objective.

**ATPase measurements**

Purified heimActin was diluted to a final concentration of 1 μM using actin polymerization buffer (20 mM Tris-HCl 8.0, 2 mM $MgCl_2$, 50 mM KCl) supplemented with 1 mM ATP. Reaction occurred at different temperatures (20 -70 $^{o}$C) for 30 min. Phosphate production was measured using a $P_i$ColorLock Gold Colorimetric Assay kit (Innova Biosciences) according to the manufacturer's instructions. Absorbance was measured at 635 nm on a PerkinElmer EnSpire microplate reader.

**Isothermal titration calorimetry**

ITC experiments were performed on an ITC200 system (MicroCal). Samples were first dialyzed against 50 mM $NaPO_4$ pH 7.5, 50 mM NaCl. Purified recombinant protein (280 µl, approx. 0.5 to 0.1 mM) were placed in the cell and titrated with 20 injections of 0.4 µl of ligand (1.0 mM polyproline (VASP)) with 1-2 min between each injection while stirring at 1500 RPM. Experiments were done at 25 °C. Data analyses were carried out using Origin 5.0 (MicroCal) provided by the manufacturer.

**Fluorescence of pyrene–actin interaction**

Pyrene-labeled rabbit actin (Cytoskeleton, Inc.) polymerization assay was performed in a 96-well, black, flat bottom plate, (Corning, Nunc). Pyrene-rabbit actin (2 µM) in G-buffer (2 mM Tris-HCl pH 8.0, 0.5 mM ATP, 0.5 mM DTT, 0.2 mM $CaCl_2$), alone or in the presence of different concentration of heimProfilin or ΔN-heimProfilin (20 µM – 280 µM) were used. The reaction was initiated with the addition of a 10x polymerization buffer (100 mM KCl and 1 mM $MgCl_2$). The increase in fluorescence intensity due to polymerization was measured at the excitation and emission wavelength of 365 and 410 nm, respectively by using the Fluoroskan Ascent FL spectrofluorometer (Thermo Scientific). The fluorescence intensity (a.u) was plotted against time using GraphPad prism.

**Sedimentation assay.**

Actin polymerization and its binding and regulation by profilin was determined by sedimentation assays. Two different buffers were used thus; Polymerization buffer 1: For ΔC-heimActin (2 mM Tris-HCl pH 7.5, 100 mM KCl, 1 mM ATP, 2 mM $MgCl_2$, 10 mM imidazol) and Polymerization buffer 2: For heimActin (20 mM Tris-HCl pH 8.0, 250 mM KCl, 1 mM ATP, 4 mM $MgCl_2$). Polymerization was initiated by diluting actin (5 µM) with 10x polymerization buffer and incubated for 2 h at room temperature. The polymerized actin filaments were pelleted by ultracentrifugation at 150,000 RCF for 1 h at 4°C using the TLA-55 rotor in Optima MAX-XP ultracentrifuge (Beckman-Coulter). The pellet was carefully separated and resuspended in the same volume as the supernatant and analyzed by SDS-PAGE. To measure the effect of heimProfilin on heimActin polymerization dynamics, 5 μM heimActin were mixed with different concentrations (0 μM, 1 μM, 5 μM, 10 μM and 20 μM) of heimProfilin or ΔN-heimProfilin and diluted with 10x buffer to initiate the polymerization. The reaction mixture was incubated for 2 h at room temperature and the filamentous actin was pelleted by ultracentrifugation at 150,000 RCF for 1 h.

**Supplementary Figure 1. Schematic of profilin and sequence alignment showing binding residues. a,** Schematic of the structure of heimProfilin. Polyproline binding residues are depicted as green sticks. **b**, An overlay of the heimProfilin (grey) and human profilin: PDB code 1pfl (blue). The position of the N and C helices of the human profilin are marked. **c**, Sequence alignment of heimProfilin (Heimd LC3) in this study and human profilin. The residues responsible for actin binding are shown in blue for the human profilin and green for heimProfilin. Also, shown in this alignment are the residues responsible for polyproline binding for human profilin (black arrows) and heimProfilin (red arrows).

**Supplementary Figure 2. Overlay of 15N-1H TROSY HSQC spectra of heimprofilin and ΔN-heimProfilin.** Shown in red is the spectrum of heimProfilin and in blue is the spectrum of ΔN-heimProfilin. The resonances of most residues overlay to each other from both proteins indicating little or no structural change. Few of the resonances in the vicinity of the deleted N-terminus experience small chemical shift change.

**Supplementary Figure 3. Purification of Heimdallarchaeota actin and profilin, rabbit actin sedimentation a,** SDS-PAGE of the purified proteins. **b, and c,** Polymerization buffer screening for the short and long actin. Both supernatant (s) and pellet (p) after ultracentrifuge were loaded onto SDS-PAGE to select the optimal buffer for polymerization. ΔC-heimActin showed the optimal polymerization in buffer 1 (2 mM Tris-HCl pH 7.5, 100 mM KCl, 1 mM ATP, 2 mM $MgCl_2$, 10 mM imidazol) while buffer 3 (20 mM Tris-HCl pH 8.0, 250 mM KCl, 1 mM ATP, 4 mM $MgCl_2$) was optimal for long actin. **d, and e,** The effect on polymerization of rabbit actin when binding with heimProfilin or ΔN-heimProfilin were monitored. The representative gels showed that heimProfilin bind with rabbit actin and delayed the polymerization, whereas no significant effect on polymerization was observed in the presence of ΔN-heimProfilin as for rabbit actin mostly appeared in pellet (p).

**Supplementary Figure 4. Nuclear magnetic resonance binding interaction of heimdallarchaeota actin and profilin.** Schematic of the structure of heimProfilin showing interacting regions for heimActin in heimProfilin **(a)** and for ΔN-heimProfilin mapped onto heimProfilin **(b)**. Expansions of overlay $^{15}$N-$^{1}$H HSQC spectra showing chemical shift changes for some of the residues for heimProfilin **(c)** and ΔN-heimProfilin with increasing concentrations of heimActin **(d).** Overlay of $^{15}$N-$^{1}$H HSQC spectra for heimProfilin **(d)** and ΔN-heimProfilin with increasing concentrations of heimActin **(e).** Note not all residues experience chemical shift changes as indicated with the arrows in e) and f).

**Supplementary Figure 5. Nuclear magnetic resonance binding interaction of heimdallarchaeota; ΔC-heimActin and ΔN-heimProfilin.** Overlay of $^{15}$N-$^{1}$H HSQC spectra of free ΔN-heimProfilin with increasing concentrations of ΔC-heimActin**.**

**Supplementary Figure 6. Sequence alignment of different profilins showing the N-terminal extension.** Protein blast search was made on NCBI and using ID OLS22855.1 (Heimdall LC3 Profilin) as input sequence. The retrieved profilins sequences were then aligned in Clustal Omega sequence alignment software. Human profilin (AAH57828.1) was also added before the alignment was made. The previous start positions are marked in red. The N-extension can be seen for a number of amino acids ranging from 5 -22 residues in length.

**Supplementary Figure 7. N-terminal extension is also present in other Asgard archaea.** Schematics of heimProfilin **(a),** and thorProfilin **(b),** showing the N-terminal extensions colored green and magenta respectively. The Thorarchaeota profilin was modeled in RaptorX while the heimProfilin 3D coordinates was determined by NMR spectroscopy in this study. The respective profilins have been reoriented to show the N-terminal extensions more clearly.

**Supplementary Figure 8. Summary of the overall interactions and schematic representation of the perceived regulatory mechanism of heimProfilin.**

Tabular summary of the interactions probed in this study **a).** The strength of the interactions have been colored coded. Red for strong, black for moderate/weak and grey-white for no interaction. **b)** HeimProfilin is depicted in cartoon with helices in red, strands in blue and loops in yellow. Possible modifications are shown in green. **i,** modification of the N-terminal loop or interaction with third-party proteins causes the heimProfilin protein to behave similarly to ΔN-heimProfilin. This prevents actin interactions but allows for $PIP_2$ and polyproline interactions. **ii**, Concurring or subsequent demodification would flip heimProfilin to an actin binding form, allowing for actin polymerization regulation. **iv**, Modification or interaction with third-party proteins would then be able to reset profilin to the first step of the cycle

**Supplementary Table 1.** Nuclear magnetic resonance spectroscopy structural statistics

| | Heimdallarchaeota profilin |
|---|---|
| **NMR distance and dihedral restraints** | |
| **Distance restraints** | |
| Total NOEs | 1000 |
| Intra-residue | 91 |
| Sequential ($\|i-j\| = 1$) | 287 |
| Medium-range ($1 < \|i-j\| < 5$) | 590 |
| Long-range ($\|i-j\| >5$) | 361 |
| **Total dihedral angle restraints** | |
| $^3J_{HN\alpha}$ scalar couplings | 83 |
| $^{13}C\alpha$ chemical shifts | 208 |
| **Structure statistics** | |
| Violations | |

| | |
|---|---|
| Distance constraints (>0.5 Å) | 0 |
| Dihedral angle constraints (>5°) | 0 |
| **Deviations from idealized geometry** | |
| Bond lengths (Å) | 0 |
| Bond angles ($^{o}$) | 0 |
| Impropers ($^{o}$) | 0 |
| [a]Average pairwise r.m.s. deviation (Å) | |
| Backbone | 0.03 ± 0.02 |
| Heavy atoms | 0.04 ± 0.03 |

---


1 Haq, S. R., Survery, S., Hurtig, F., Lindas, A.-C. & Chi, C. N. NMR backbone assignment and dynamics of Profilin from Heimdallarchaeota. *bioRxiv*, 2020.2004.2020.050468, doi:10.1101/2020.04.20.050468 (2020).

2 Vranken, W. F. *et al.* The CCPN data model for NMR spectroscopy: development of a software pipeline. *Proteins* **59**, 687-696, doi:10.1002/prot.20449 (2005).

3 Guntert, P., Mumenthaler, C. & Wuthrich, K. Torsion angle dynamics for NMR structure calculation with the new program DYANA. *J Mol Biol* **273**, 283-298, doi:10.1006/jmbi.1997.1284 (1997).

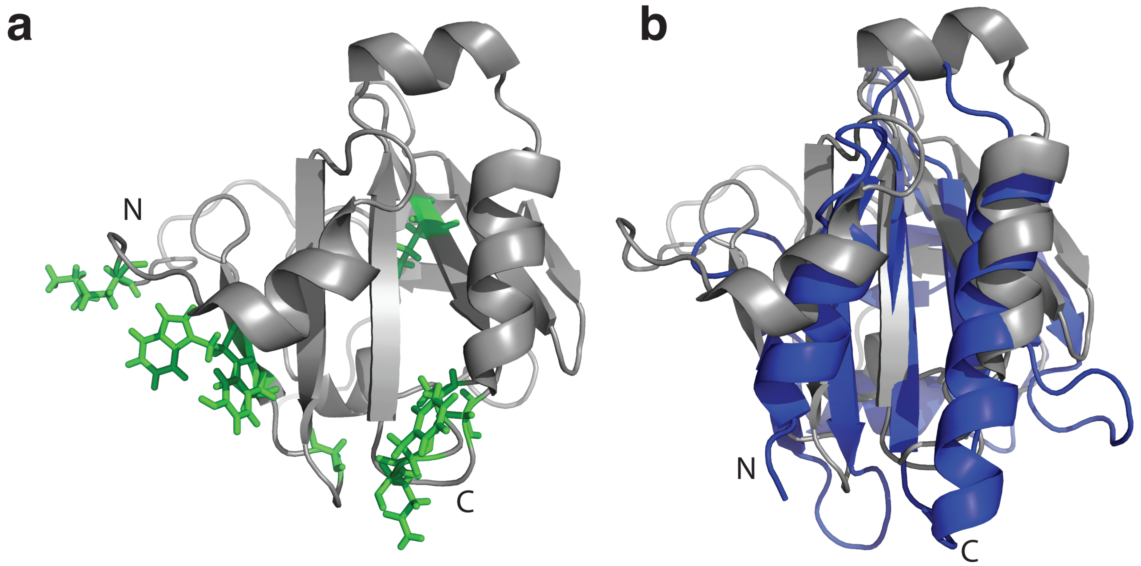


**c**

```
Human     1 ----------------------MA--GWNAYID-NLMA---DGTCQDAA-IVGYKDSPSVWAAVPGKTFVNITPAEVGVLVG--KDRSSFY---VNGLTLGGQKCSVIRDSLLQDGE
Heimdall  1 MKDGFILKKKKIPKNIFRIGNKMS---YSDDLN-SLIQ---SQYVSYVV-ILDPNG--AIYWTNNEN-W-QVN-GSEVLRQWMGSA---------PSITVAGTKFSSFRNEPG----

Human       FSMDLRTKSTGGAPTFNVTVTK----TDKTLVLLMGKEGVHGGLINKK--YEMASHLRRSQY- 140
Heimdall    --VSFVGRNMAGGG---LIILQKAP-NGY-VFLSWTSHEFLTTSGLPPLNIHAEIAMMAAKFQ 148
```

S. 1

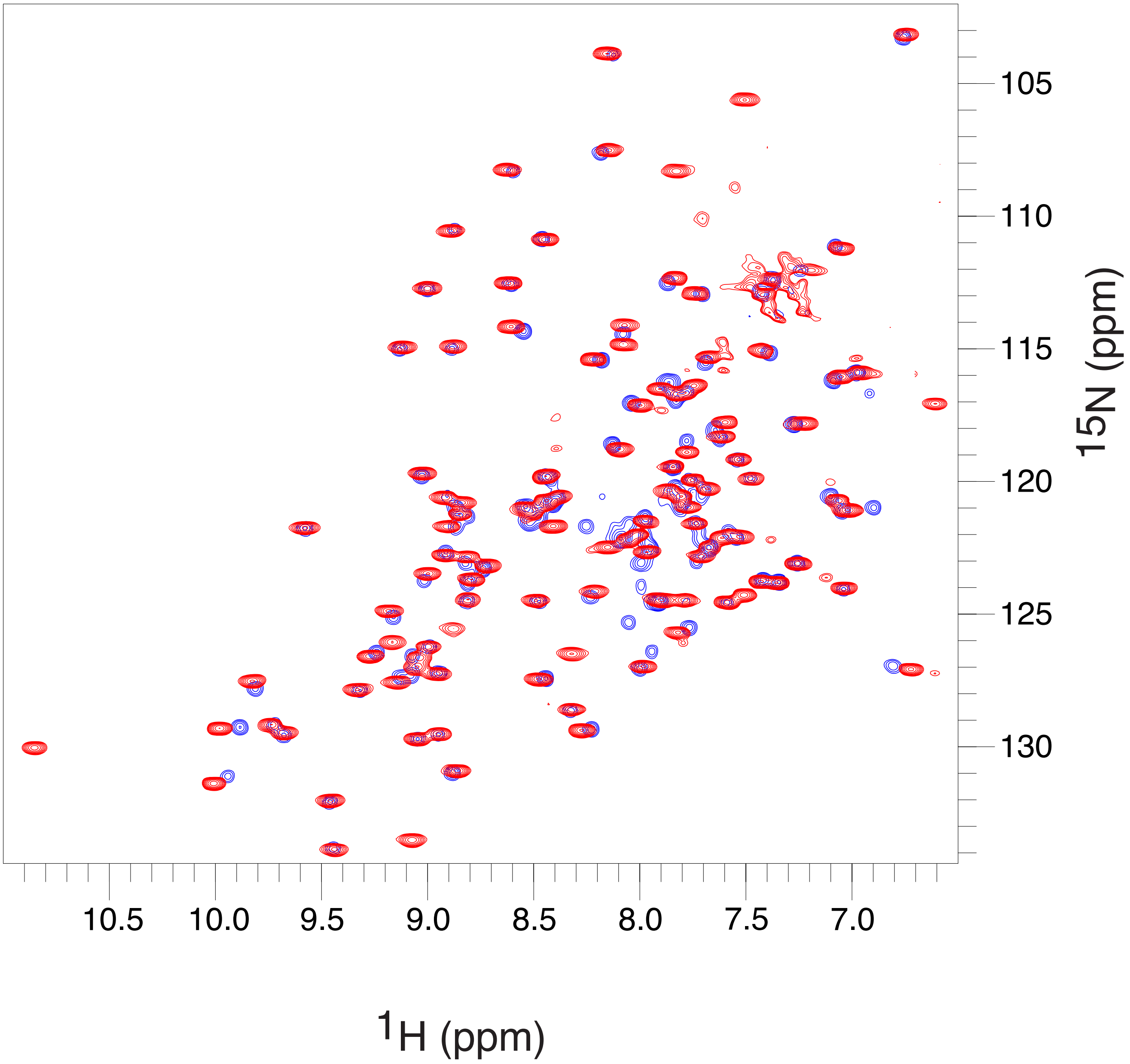


S. 2

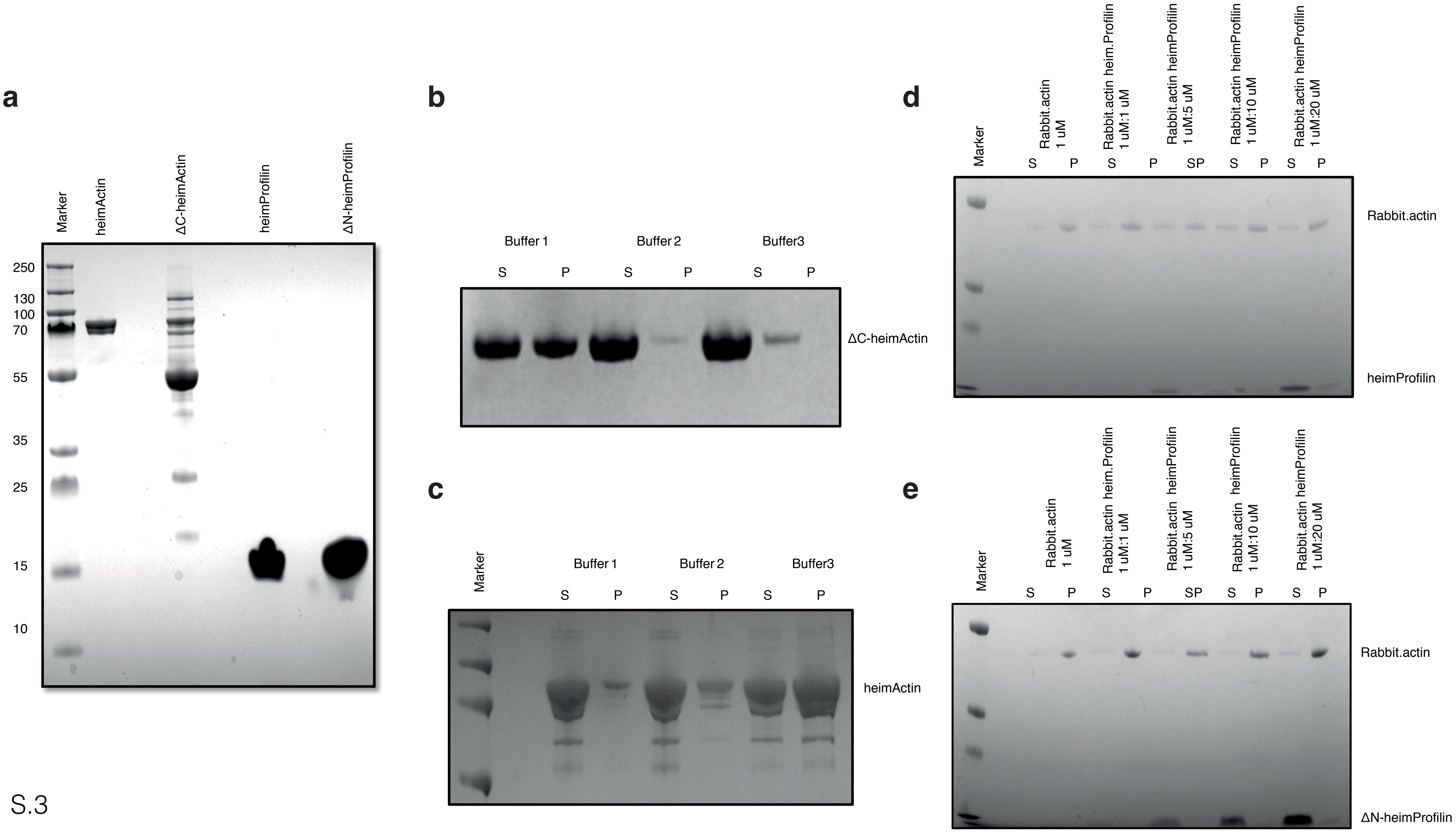
a
Marker
heimActin
ΔC-heimActin
heimProfilin
ΔN-heimProfilin
250
130
100
70
55
35
25
15
10
b
Buffer 1
Buffer 2
Buffer3
S
P
S
P
S
P
ΔC-heimActin
c
Marker
Buffer 1
Buffer 2
Buffer3
S
P
S
P
S
P
heimActin
d
Marker
Rabbit.actin 1 uM
Rabbit.actin heim.Profilin 1 uM:1 uM
Rabbit.actin heimProfilin 1 uM:5 uM
Rabbit.actin heimProfilin 1 uM:10 uM
Rabbit.actin heimProfilin 1 uM:20 uM
S
P
S
P
SP
S
P
S
P
Rabbit.actin
heimProfilin
e
Marker
Rabbit.actin 1 uM
Rabbit.actin heim.Profilin 1 uM:1 uM
Rabbit.actin heimProfilin 1 uM:5 uM
Rabbit.actin heimProfilin 1 uM:10 uM
Rabbit.actin heimProfilin 1 uM:20 uM
S
P
S
P
SP
S
P
S
P
Rabbit.actin
ΔN-heimProfilin

S.3

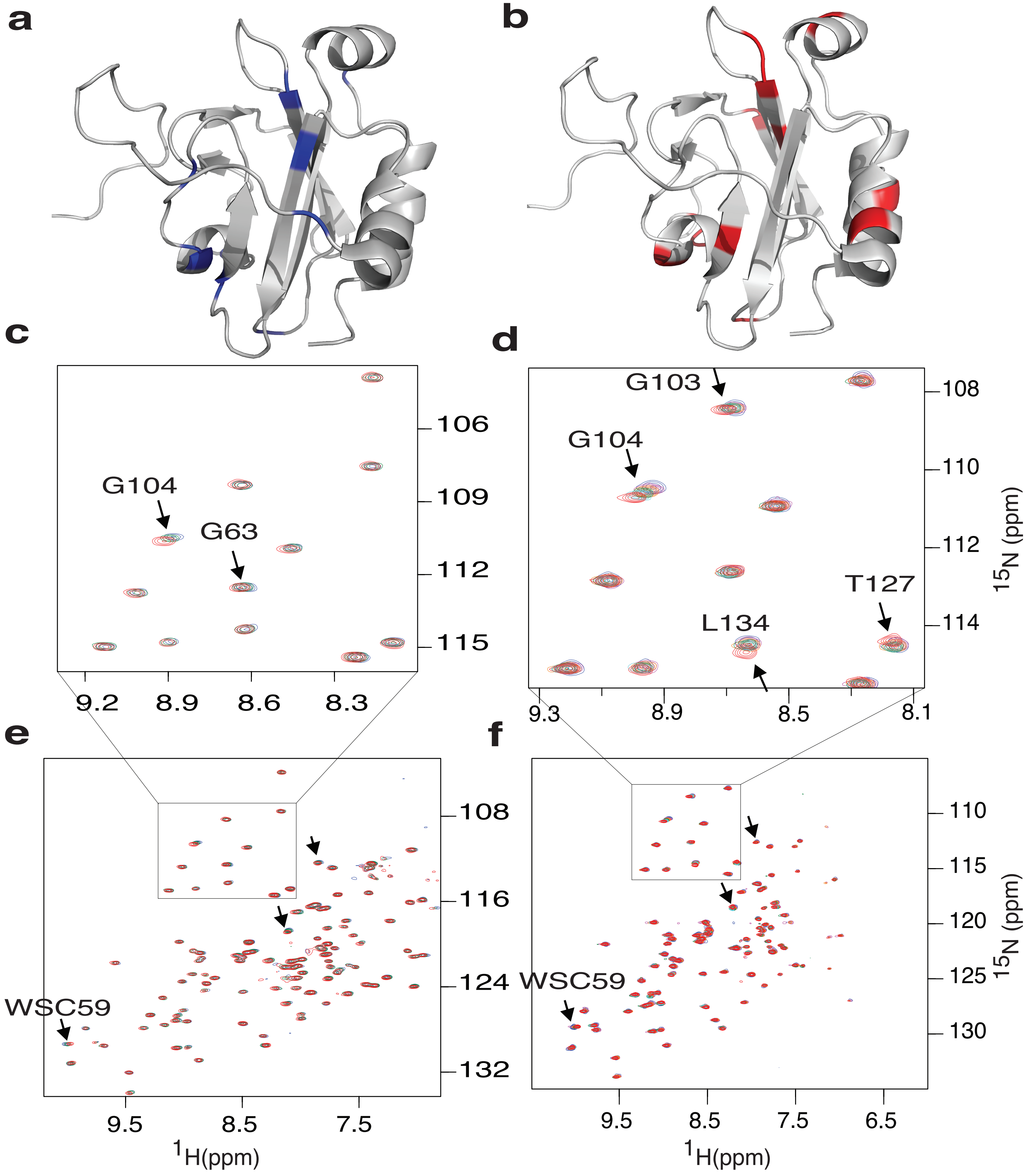
a
b
c
d
e
f
G104
G63
106
109
112
115
9.2
8.9
8.6
8.3
G103
G104
T127
L134
108
110
112
114
$^{15}$N (ppm)
9.3
8.9
8.5
8.1
WSC59
108
116
124
132
9.5
8.5
7.5
$^{1}$H(ppm)
WSC59
110
115
120
125
130
$^{15}$N (ppm)
9.5
8.5
7.5
6.5
$^{1}$H(ppm)

S.4

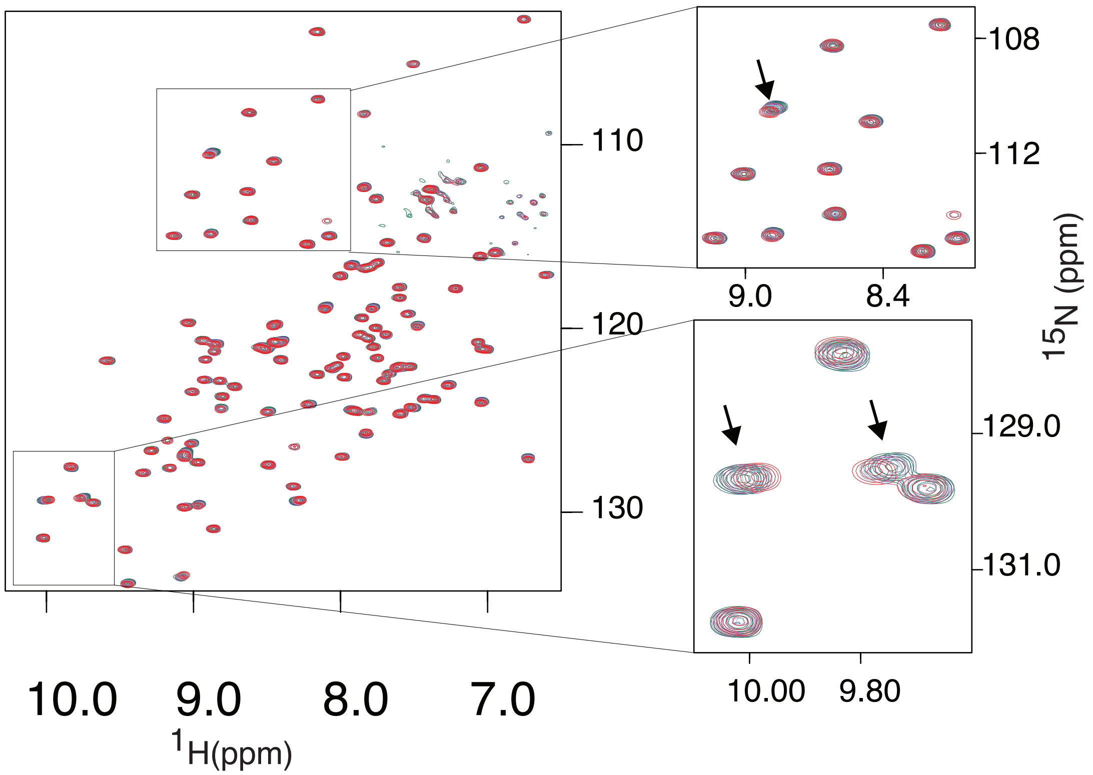

110
120
130
10.0
9.0
8.0
7.0
$^{1}$H(ppm)
108
112
9.0
8.4
$^{15}$N (ppm)
129.0
131.0
10.00
9.80


S.5

```
GenBank: ID      --------------------------Profilin Sequence-------------------

AAH57828.1       --------------------MAGWNAY-I-DNLMADGTCQDAAIVGYKDSPSVWAAV--- 35
HDH89712.1       --------------------MSEEAINIISELMTSNPTITALAIVS-SNDQVLWQTP--- 36
RLI70375.1       --------------------MSFEA---EAARLQREGIVGALALIG-ADGQVYWQTP--- 33
TFG12995.1       -MCGYNLGTEWISLRSKLEMSDPVMEAYTQAYEASNQMVQAFGVIK-ADGSVLWQSN--- 55
TFG09823.1       ------------MNRGEFRMADPVMEAYTQAYNDSNEMVQAFGIIT-TSGNVLWQSD--- 44
TFG30347.1       -------------------MSDPVMDAYTMAYNSSGNMVQAFGVIT-ESGHVLWQSN--- 37
TFF94849.1       -------------------MSDPVMDAYTKAYNSSNNMVQAFGVIT-EGGQVLWQSN--- 37
RLI55859.1       --------MVDPQERGEFTMSDPVMDAYTQAYNSSGNMVQAFGVIT-EGGQVLWQSN--- 48
PWI46981.1       -----------------------MPYRQPLEKLLEEKKVLGVVIFD-PSGNDWWHTGSFP 36
OLS22855.1       MKDGFILKKKKIPKNIFRIGNKEMSYSDDLNSLIQSQYVSYVVILD-PNGAIYWTNN--- 56
PWI48213.1       -----------------------MSYQDDMNALIQNGYVSIVTILD-PNGAPYWTNQ--- 33
                                                            :.   ..   *
GenBank: ID      --------------------------Profilin Sequence-------------------

AAH57828.1       ---------PGKTFVNIT-PAEVGVLVGKDRSS----------FYV-NGLT-------LG 67
HDH89712.1       ---SWNIQ--GSRIISAWMNREPSVVVQGVKYSTFQAEPE--KFIAKN----------MS 79
RLI70375.1       ---NWQVD--GRSIIEAWTKKAPWVVVQGVKYSTIDITNN--RLICTN----------VQ 76
TFG12995.1       ---NWDLSADAQQLISAVNNQSASVKQNDVKYSTIRTSPE--SLVARN----------VQ 100
TFG09823.1       ---NWDLTEDAAALGSAVKSSATSVVQNQVKYSTLRTTPE--SLVARS----------VS 89
TFG30347.1       ---NWDLTADASALMSAVKSRSPAITQNNVRYSTMRSTPE--SLVARN----------VQ 82
TFF94849.1       ---NWDLTADAAGLMAAVKSRSASVIQNQVKYSTMRTTPE--SLVARN----------VQ 82
RLI55859.1       ---NWDLTADAANLIAAVRSRAASVVQNQVKYSTMRTTPE--SLVAKS----------IG 93
PWI46981.1       QANDRLLD--GYYLIHEWVTYPASAEIAGVKYLTLLNAYPD-YWLLTN----------MQ 83
OLS22855.1       --ENWQVN--GSEVLRQWMGSAPSITVAGTKFSSFRNEPGV-SFVGRN----------MA 101
PWI48213.1       --PEWQVD--GPALLSSWQNKEPGVNIAGVRYSSMVNDWEVGNYVARN----------VG 79
                     .                          : .:         :  .          :

GenBank: ID      --------------------------Profilin Sequence-------------------

AAH57828.1       GQKCSVIRDSLLQDGEFSMDLRTKSTGGAPTFNVTVTKTDKTLVLLMGKEGVHGGLINKK 127
HDH89712.1       GYGYVVATPLT--NELFLVAYVTPE--------------------------GDPKAAFVD 111
RLI70375.1       QQGHVIIAKTPF-WQGFVMAWCSPA--------------------------NPVRLIYVD 109
TFG12995.1       GNGTLILARIE--DDKWVVAWASAD--------------------------AAPDSVYVD 132
TFG09823.1       GNGILVLAKIE--GEKFVVAWADPN--------------------------ASPDGIYVD 121
TFG30347.1       GNGILILAKIE--GEKWVVAWVAPD--------------------------AQPDGVYVD 114
TFF94849.1       GSGILVLTKIDASSDRWVVAWADPN--------------------------AQPDGIYVD 116
RLI55859.1       GNGILILTKVDAASDRWVVAWADPN--------------------------AQPDGIYVD 127
PWI46981.1       GQGSLILQKAT--NGFYFFCYLDES--------------------------IDPPDIQKE 115
OLS22855.1       GGGLIILQKAP--NGYVFLSWTSHEFL--------------------TTSGLPPLNIHAE 139
PWI48213.1       GAGIICLVRAP--NNYYFLTWTPGD------------------------VQIPSINVHGE 113
                 .                 .                                        .
GenBank: ID      -Profilin Sequence------

AAH57828.1       CYEMASHLRRSQY         140
HDH89712.1       LYNAAIRIRKVI-         123
RLI70375.1       VAKLASTVKPG--         120
TFG12995.1       IDRAAKALKGKI-         144
TFG09823.1       VDRAAKALKGKI-         133
TFF91518.1       VDRAAKALKGKI-         126
TFG30347.1       VDRAAKTLKGRI-         126
TFF94849.1       VDRAAKTLKGKI-         128
RLI55859.1       VDRAAKTLRGKI-         139
PWI46981.1       VKVIAELFAQGV-         127
OLS22855.1       IAMMAAKFQ----         148
PWI48213.1       VAKMAIKFQ----         122
```

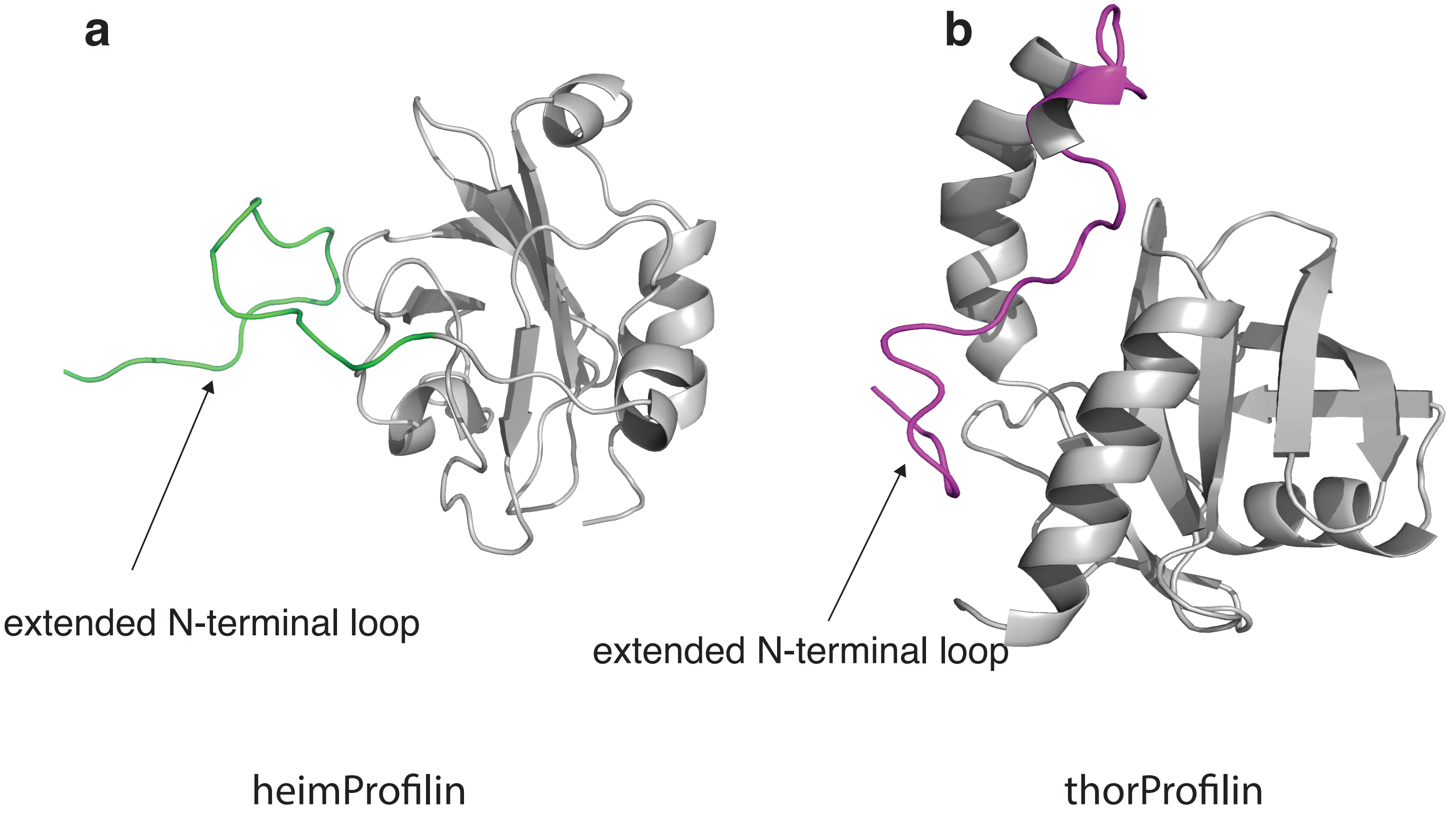
a
b
extended N-terminal loop
extended N-terminal loop
heimProfilin
thorProfilin

S.7

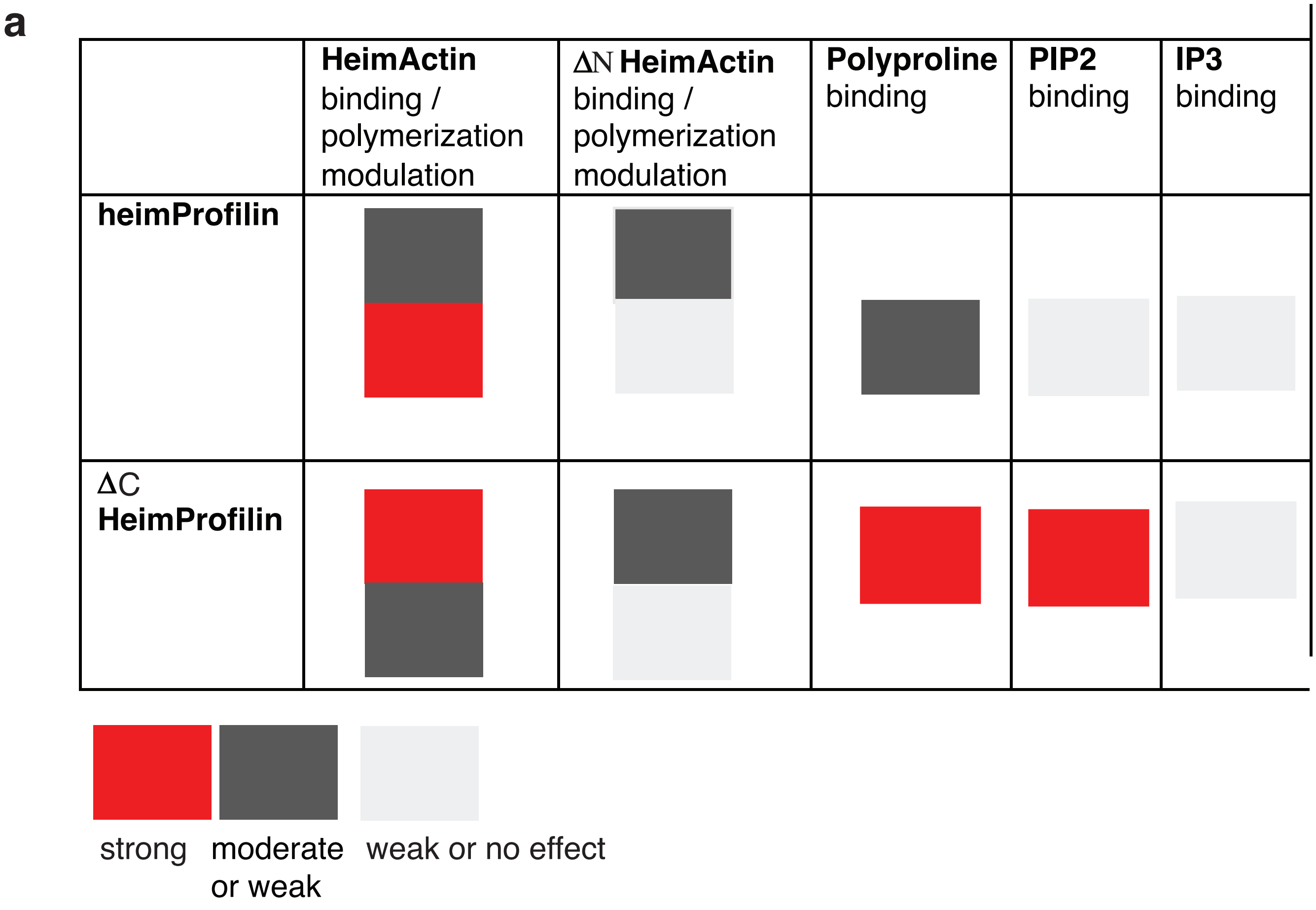
a
HeimActin binding / polymerization modulation
ΔN HeimActin binding / polymerization modulation
Polyproline binding
PIP2 binding
IP3 binding
heimProfilin
ΔC HeimProfilin
strong
moderate or weak
weak or no effect

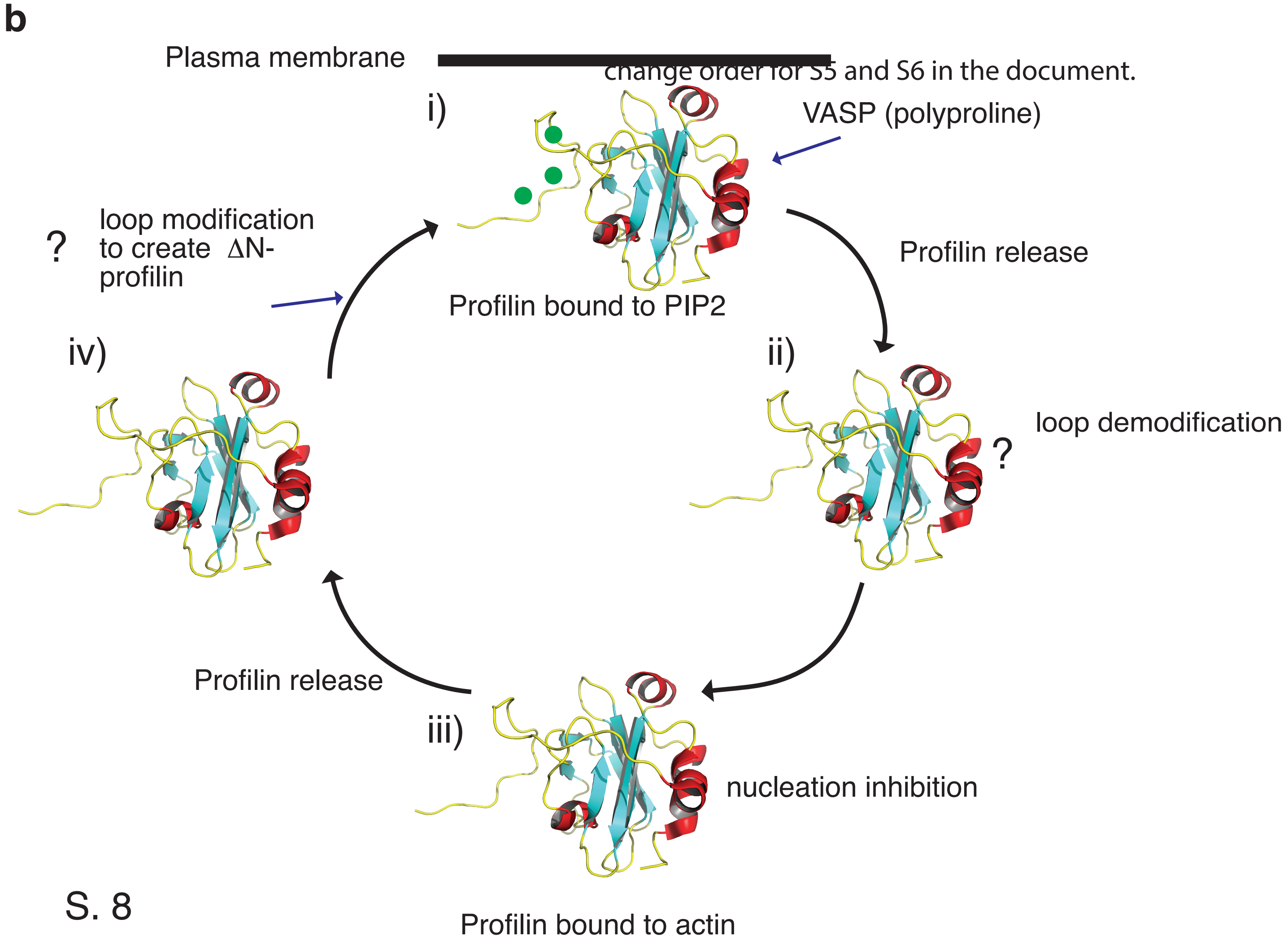
b
Plasma membrane
change order for S5 and S6 in the document.
i)
VASP (polyproline)
Profilin bound to PIP2
Profilin release
ii)
loop demodification
?
iii)
nucleation inhibition
Profilin bound to actin
Profilin release
iv)
?
loop modification to create ΔN-profilin

S. 8
